# Microsomal triglyceride transfer protein is necessary to maintain lipid homeostasis and retinal function

**DOI:** 10.1101/2023.12.06.570418

**Authors:** Catharina R. Grubaugh, Anuradha Dhingra, Binu Prakash, Diego Montenegro, Janet R. Sparrow, Lauren L. Daniele, Christine A. Curcio, Brent A. Bell, M. Mahmood Hussain, Kathleen Boesze-Battaglia

**Author notes:** Corresponding author: Dr. Kathleen Boesze-Battaglia University of Pennsylvania, Department of Basic and Translational Sciences 240 South 40th Street, Philadelphia, PA 19014.

## Abstract

Lipid processing by the retinal pigment epithelium (RPE) is necessary to maintain retinal health and function. Dysregulation of retinal lipid homeostasis due to normal aging or to age-related disease triggers lipid accumulation within the RPE, on Bruch’s membrane (BrM), and in the subretinal space. In its role as a hub for lipid trafficking into and out of the neural retina, the RPE packages a significant amount of lipid into lipid droplets for storage and into apolipoprotein B (apoB)-containing lipoproteins (Blps) for export. Microsomal triglyceride transfer protein (MTP), encoded by the *MTTP* gene, is essential for Blp assembly. Herein we test the hypothesis that MTP expression in the RPE is essential to maintain lipid balance and retinal function using the newly generated *RPEΔMttp* mouse model. Using non-invasive ocular imaging, electroretinography, and histochemical and biochemical analyses we show that genetic deletion of *Mttp* from the RPE results in intracellular lipid accumulation, increased photoreceptor –associated cholesterol deposits and photoreceptor cell death, and loss of rod but not cone function. RPE-specific ablation of Mttp had no significant effect on plasma lipids and lipoproteins. While, apoB was decreased in the RPE, ocular retinoid concentrations remained unchanged. Thus suggesting that RPE MTP is critical for Blp synthesis and assembly but not directly involved in ocular retinoid and plasma lipoprotein metabolism. These studies demonstrate that RPE-specific MTP expression is necessary to establish and maintain retinal lipid homeostasis and visual function.

## Introduction

Visual function depends on the intimate structural, functional and metabolic interactions amongst the retinal pigment epithelium (RPE), neural retina (NR) and choriocapillaris (1–3). A growing body of research highlights the importance of local pathways (i.e., those operating within the eye) in the regulation of retinal lipids. In particular, the RPE is a central hub for lipid processing within the eye and acts as a gatekeeper for lipid transport into and out of the NR (1, 4–7). In this role, the RPE ingests fragments of outer segments (OS) of photoreceptors (PR) in a process historically called “phagocytosis,” but more accurately referred to as “trogocytosis,” or “nibbling,” (8). In the human retina, Volland et al. (2015) estimate that each RPE cell ingests and processes ∼0.30 pmol of phospholipid per RPE cell (9). In addition to the processing of lipid rich-OSs, the RPE synthesizes lipids *de novo*, and takes up plasma lipids from the choroidal circulation through its basal membrane. The RPE also transfers lipids from these three sources to the NR through its apical membrane (3), with the recycling of the polyunsaturated fatty acids, docosahexaenoic acid (DHA) essential for function (6, 9–11).

Given that lipid homeostasis between the RPE and NR is critical to retinal health and, thereby, visual function, the RPE must employ mechanisms to prevent steatosis due to its considerable lipid intake. In this regard, the RPE oxidizes lipids and generates β-hydroxybutyrate (βHB), which it releases apically, providing the PR with metabolic substrate (12, 13). To facilitate lipid export to the NR and into the systemic circulation, the RPE packages excess lipids into apolipoprotein β-containing lipoproteins (Blps) (14, 15). These Blps consist of a lipid core, rich in esterified cholesterol (EC) and triglycerides (TG), enclosed in a layer of phospholipids and apolipoproteins, including apolipoprotein-β (APOB). Such an arrangement facilitates *en mass* transport of hydrophobic lipids through the aqueous environment (16, 17). Therefore, the RPE plays an essential homeostatic role for processing and bidirectional export of lipids, including cholesterol, for the entire RPE/NR/choroid complex (1–3, 5, 18).

Numerous studies encompassing *in vitro* systems, *in vivo* models, and patient data have indicated that defects in lipid processing pathways and metabolic dysregulation in the RPE contribute to aging, retinitis pigmentosa and age-related retinal disease (14–16, 19–21). Mouse models of defective phagosome maturation and of dysfunctional lipid processing pathways, have documented RPE steatosis-like lipoidal degeneration (22, 23), loss of RPE and/or PR function, and inflammation (5, 6, 19). For instance, RPE-specific ablation of the ATP-dependent transporter ABCA1, which exports cholesterol to the NR, (3, 10, 11) results in RPE lipid accumulation, loss of RPE and PR function, and retinal degeneration (2). In AMD, a critical role of lipids is highlighted by the presence of lipids in pathognomonic extracellular deposits called drusen (14, 19). The role of RPE-specific lipid processing pathways in health and disease, such as in the development of sub-retinal and sub-RPE deposits, remains to fully explored.

Understanding how local lipoprotein production and secretion forestalls steatosis in the outer retina is critical for identifying therapeutic targets for retina–specific lipid accumulation. The objective of the current study is to assess the role of RPE-specific MTP expression in maintaining retinal lipid homoeostasis. We hypothesized that Blp assembly in the RPE, a local mechanism to regulate lipid processing and transport, is necessary to maintain retinal health and function. To test this hypothesis, we generated the *RPEΔMttp* mouse model that has an RPE-specific deficiency of MTP. In the absence of MTP, RPE of *RPEΔMttp* mice accumulated large amounts of lipids, suggesting that a major function of MTP in the RPE might be to prevent lipid accumulation by exporting lipids out of the eye and into the choroidal circulation. Our results highlight the essential role of MTP-mediated Blp assembly in the RPE and demonstrate the pathogenic consequences of decreased lipid transport via Blps assembled in the RPE.

## Materials and Methods

### Animals

Mice homozygous for the floxed *Mttp* allele (*Mttp^flox/flox^*) on a C57Bl6/J background (24–28)were crossed with a transgenic line that expresses Cre recombinase under control of the human bestrophin 1 (*BEST1*) promoter (C57BL/6-Tg(BEST1-cre)1Jdun/J, referred to as Best1-Cre; Jax stock no. 017557) (29) which were purchased from Jackson Laboratories (Bar Harbor, ME, USA). Two-generation crosses were used to produce mice that were *Mttp^flox/flox^*and Cre transgenic (*Mttpf^lox/flox^; BEST1-Cre^+/−^*, hereafter referred to as *RPEΔMttp*). In all of our experiments, *RPEΔMttp* mice were compared to age-matched controls, which were homozygous for the wild-type *Mttp* allele and hemizygous for the transgenic Cre allele (*Mttp^wt/wt^*and *BEST1-Cre^+/−^*, hereafter referred to as Controls). Mouse lines were confirmed to be free of retinal-degeneration related mutations, rd8 and rd10, by genotyping (Transnetyx, Cordova, TN,USA) (30, 31). The mice were genotyped for *Mttp* and *Cre* by PCR on tail genomic DNA using the following primers:

*Mttp* primers:

MTP forward: 5’ – GAGCCTGTTGAGCAAGTACAAG – 3’
MTP reverse: 5’ – GGCAGCAGGACAGAGACAC – 3’

*Cre* primers

Best1-Cre F: 5’ – TCGATGCAACGAGTGATGAGG – 3’
Best1-Cre R: 5’ – GGCCCAAATGTTGCTGGATAG – 3’

Maintenance of mouse colonies and all experiments involving animals were as described previously (22). Mice were housed under a 12-hr light/dark cycle and fed *ad libitum* (22) with both female and male mice used in these studies. All procedures involving animals were approved by the Institutional Animal Care and Use Committee at the University of Pennsylvania and were in accordance with the Association for Research in Vision and Ophthalmology (ARVO) guidelines for use of animals in research.

### Antibodies

Commercially available primary antibodies used are shown in Table 1. Secondary antibodies used are: goat anti-Mouse and goat anti-rabbit horseradish peroxidase (HRP)-conjugated antibodies (Invitrogen, Waltham, MA, USA) for Western blotting, and donkey anti-mouse, anti-rabbit, anti-goat IgG Alexa Fluor 488/ 594/ 647 conjugates (Invitrogen) for immunostaining experiments.

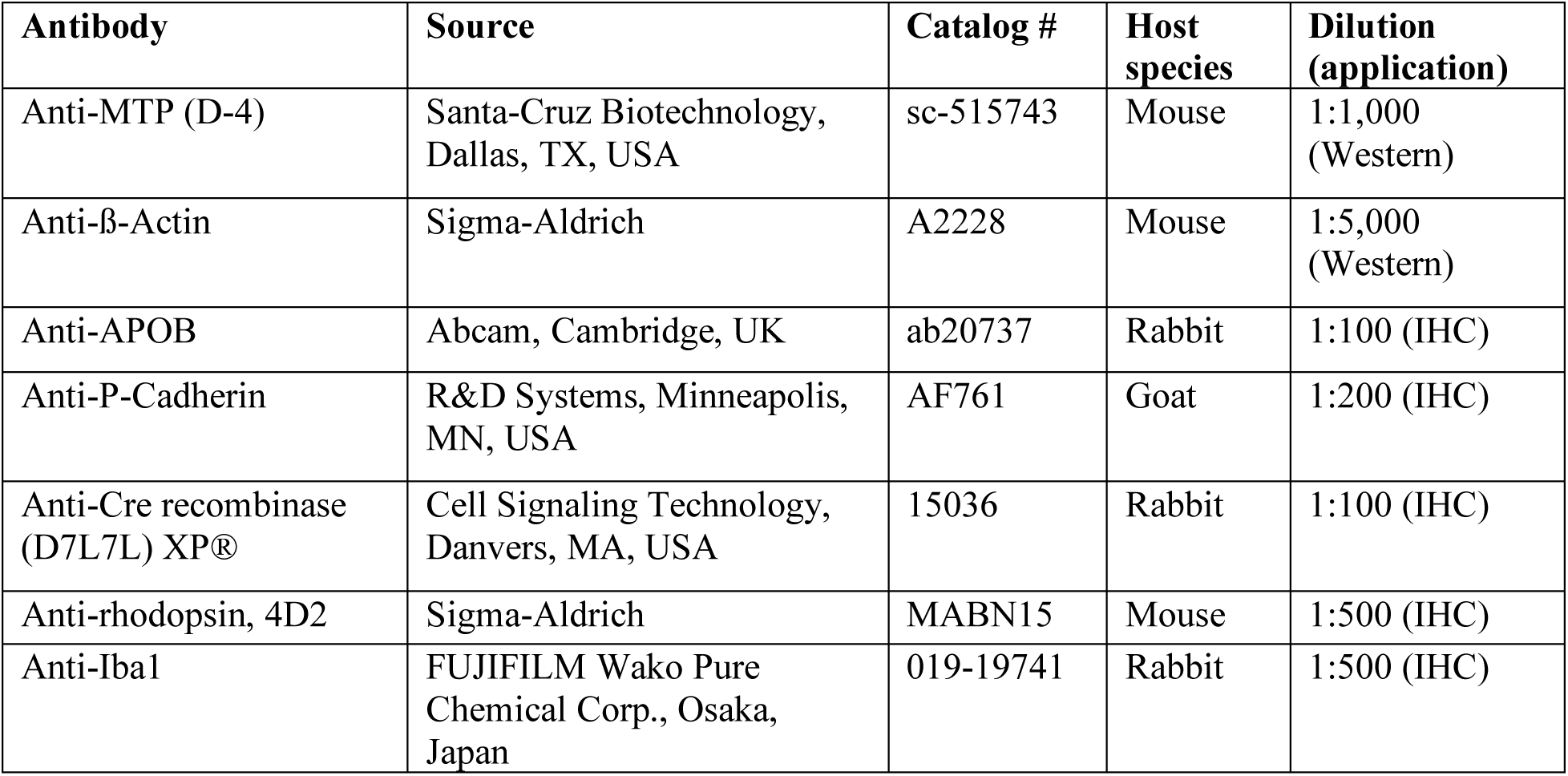

| Antibody | Source | Catalog # | Host species | Dilution (application) |
| --- | --- | --- | --- | --- |
| Anti-MTP (D-4) | Santa-Cruz Biotechnology, Dallas, TX, USA | sc-515743 | Mouse | 1:1,000 (Western) |
| Anti-β-Actin | Sigma-Aldrich | A2228 | Mouse | 1:5,000 (Western) |
| Anti-APOB | Abcam, Cambridge, UK | ab20737 | Rabbit | 1:100 (IHC) |
| Anti-P-Cadherin | R&D Systems, Minneapolis, MN, USA | AF761 | Goat | 1:200 (IHC) |
| Anti-Cre recombinase (D7L7L) XP® | Cell Signaling Technology, Danvers, MA, USA | 15036 | Rabbit | 1:100 (IHC) |
| Anti-rhodopsin, 4D2 | Sigma-Aldrich | MABN15 | Mouse | 1:500 (IHC) |
| Anti-Iba1 | FUJIFILM Wako Pure Chemical Corp., Osaka, Japan | 019-19741 | Rabbit | 1:500 (IHC) |

### BaseScope *In situ* Hybridization

*Mttp* RNA *in situ* hybridization was performed on fixed frozen sections using the BaseScope 2.5 HD assay kit (Advanced Cell Diagnostics, Newark, CA) (32, 33). A custom-designed BaseScope probe targeting bases 855-989 of the mouse *Mttp* transcript, NM_008642.3, which span exon 5 and part of exon 6 was used for the assay (BaseScope™ Probe-BA-Mm-Mttp-3zz-st-C1; Cat # 1217461-C1; Advanced Cell Diagnostics, Minneapolis, Minnesota, USA). Briefly, mouse retinal cryosections (10 µm thick) from ∼12 month-old *RPEΔMttp* and control mice retinas were post-fixed in 4% paraformaldehyde (PFA) for 60 mins at 4°C, followed by dehydration in ethanol, bleaching (10% H2O2 at 60°C), target retrieval (5 min), dehydration, and Protease III treatment (15 min at 40°C). The sections were incubated with the BaseScope probe for 2 h at 40°C, followed by Hybridize BaseScope™ v2 AMP steps 1-8 and signal detection per manufacturer’s instructions (BaseScope™ Detection Reagent Kit v2 – RED; Document # 323900-USM; Advanced Cell Diagnostics). The sections were counterstained with Hoechst 33258 (1:10,000) and mounted with Prolong Gold Antifade Mountant (P36930, Molecular Probes, Waltham, MA, USA).

### Immunoblotting

RPE/choroid (RPE/Ch) lysates were prepared in RIPA buffer with 1% protease inhibitor mixture (P8340, Sigma-Aldrich, Burlington, MA, USA) and 1% phosphatase inhibitor cocktail 2 (1862495, Thermo Fisher Scientific, Waltham, MA, USA). Lysates of (20-30 μg of protein) were separated on NuPAGE 4-12% Bis-Tris gels (Invitrogen) under reducing conditions and transferred to PVDF membranes (Millipore, Burlington, MA, USA). Membranes were blocked with 5% milk in Tris-buffered saline (TBS), 0.1% Tween-20 (TBST) for 1h at room temperature and incubated with primary antibodies overnight at 4°C. Membranes were washed and incubated with goat anti-mouse (1:2,500) HRP-conjugated secondary antibodies for 1h at room temperature. Blots were developed using ECL SuperSignal® West Femto extended duration substrate (Thermo Fisher Scientific) and captured on Odyssey Fc (LI-COR Biosciences, Lincoln, NE, USA) and quantified as previously described (12).

### Immunostaining

Immunostaining was performed on mouse retinal cryosections or RPE/Ch flat mounts as previously described (22). For retinal cryosections, mice were euthanized, and the eyes were enucleated, incised at the ora serrata and fixed in 4% PFA in PBS (pH 7.4) for 18 h at 4 °C followed by washes with PBS to remove fixative. The eyes were then cryoprotected in 30% sucrose, embedded in OCT, and stored at −80°C. Cryosections were prepared by radially sectioning the blocks at 10 µm thickness. RPE/Ch flat mounts were prepared by separating the retina from the RPE followed by fixation in 4% PFA for 30 min at room temperature. The sections or flat mounts were permeabilized and blocked with 5% BSA in PBST (0.2% Triton X-100 in PBS) at 37°C for 1 h, incubated with the primary antibody diluted in blocking solution (dilution factor as in Table 1) at 4°C overnight, washed three times with PBST, incubated in appropriate secondary antibodies conjugated to Alexa Fluor dyes (Invitrogen, 1:1,000) and Hoechst 33258 (1:10,000) at 37 °C for 1h and washed three times with PBST. In some experiments with cryosections, Alexa Fluor 647 Phalloidin (1 Unit/200µl) (Invitrogen; A22287) was included in the secondary antibody step.

### Lipid Staining

For cholesterol staining with Filipin, retinal cryosections were air dried, rinsed in PBS, and permeabilized in 0.5% Triton X-100 for 20 mins at 37°C followed by PBS washes. The sections were then incubated in 60 μg/ml Filipin (Sigma-Aldrich, F9765) for 2 h at room temperature in the dark, followed by PBS washes (PMC4235331) (22). The sections were counterstained with 5 µM DRAQ5 (Cell Signaling, #4084), washed in PBS, mounted in Prolong Gold Antifade and imaged. For LipidTOX™ staining, flat mounts were incubated in 1:200 dilution of HCS LipidTOX™ Red (Thermo Fisher Scientific; H34476) in PBS for 30 min at room temperature (RT) and washed in PBS. Sections were mounted in Prolong Gold Antifade Mountant and imaged.

### Confocal Imaging

Images were captured on a Nikon (Minato City, Tokyo, Japan) A1R laser scanning confocal microscope with a 10X(NA 0.45) or 20X (NA 0.75) dry objective, a PLAN APO VC 60X (NA 1.2) water objective, or a CFI60 Plan Apochromat Lambda 100X (NA 1.45) oil objective at 18 °C. Data were analyzed using Nikon Elements 5.30.03 AR software (22). For quantification of APOB staining, background subtraction was performed on the z-stack images, and the mean ROI intensity for each field was determined using the Nikon Elements software. Mean intensity values for the RPEΔMttp ( 7 fields from 2 mice/genotype) was normalized as % of control.

### APOB and Cholesterol Quantification

APOB protein levels in RPE and retinal lysates were measured using an Apo B ELISA Kit (ab230932, Abcam Cambridge, UK). Briefly, tissues were homogenized in chilled 1× Cell Extraction Buffer (5X PTR; ab193970, Abcam), incubated on ice for 20 minutes, and centrifuged at 18,000-× g for 20 minutes at 4°C. Cleared supernatant was collected, and protein concentration was measured using Micro BCA™ Protein Assay Kit (23235, Thermo Fisher Scientific). Samples were diluted to 200 µg/ml in 1x Cell Extraction Buffer, and the assay performed per the manufacturer’s specifications.

Total cholesterol (esterified and unesterified) in neural retina lysates was measured using Amplex™ Red Cholesterol Assay Kit (A12216) per the manufacturer’s instructions (Thermo-Fischer) as we have described previously (34). Intensity measurements were performed with an Infinite 200 Pro plate reader (Tecan Group Ltd., Männedorf, Switzerland) with at λex=555nm and λem=580nm.

### Electron Microscopy

*RPEΔMttp* and age-matched Best1-Cre control mice were enucleated, the anterior segment removed, and eyecups immersed in fixative (2.5% glutaraldehyde and 2% paraformaldehyde, in 0.1 M cacodylate buffer) overnight at 4°C, rinsed with sodium cacodylate buffer (pH 7.4), trimmed into ∼2 mm pieces and post-fixed in 1% osmium tetroxide. Samples were, stained with uranyl acetate, dehydrated and processed for embedding into Epon as described (35–37). Semi-thin sections (∼1μm thick) were obtained along the dorso-ventral axis, stained with toluidine blue, and imaged using Nikon Eclipse 80i microscope with 40X and 100X objectives. Ultrathin sections (60-80 nm) were imaged with a Jeol-1010 transmission electron microscope. Post-fixation, embedding, and ultrathin sectioning were performed by the University of Pennsylvania Electron Microscopy Resource Laboratory.

### *In vivo* Retinal Imaging

*RPEΔMttp* and age-matched control mice were imaged at 5 or 7 months of age using a Spectralis HRA confocal scanning laser ophthalmoscope (cSLO; Heidelberg Engineering, Inc., Franklin, MA, USA) and Bioptigen Envisu R2200 ultra-high resolution (UHR) spectral-domain optical coherence tomography system (SD-OCT; Leica Microsystems, Deerfield, IL, USA) (22). Mice were prepared for *in vivo* imaging by applying one drop of a 2:1 mixture of 1% Tropicamide HCl (Benzeneacetamide, N-ethyl-α-(hydroxymethyl)-N-(4-pyridinylmethyl) and 2.5% Phenylephrine (Akorn, Inc., Lake Forest, IL, USA) to both eyes several minutes prior to anesthesia. Anesthesia was achieved with an intraperitoneal injection of 95 mg/kg of Ketamine HCL (2-(2-chlorophenyl)-. 2-(methylamino)-cyclohexanone, Dechra Veterinary Products, Overland Park, KS, USA) and 10-11 mg/kg of Xylazine HCL (N-(2,6-dimethylphenyl)-5,6-dihydro-4H-1,3-thiazin-2-amine, Akorn, Inc.). Following general anesthesia, corneas received topical anesthesia with 1% Tetracaine (Alcon Laboratories, Inc., Ft. Worth, TX, USA) followed a few minutes later by application of Refresh Artificial Tears (Allergan, Irvine, CA, USA) in conjunction with protective eye shields to prevent corneal desiccation and media opacity development (38). Mice were then relocated to the Spectralis imaging platform for cSLO imaging.

Two cSLO images were collected using a Ultrawide Field (UWF, Heidelberg Engineering, Inc., Franklin, MA, USA) 105° lens that provided ∼3.4 mm field of view (FOV) of the mouse fundus. The first image was a dark-field (i.e. cross-polarization) infrared IR reflectance image used to center the optic disk in the real-time field of view window and train the focus on the RPE-choroid interface. Following alignment, a blue autofluorescence (BAF) image, which has a 486 nm raster scanned laser for excitation and 500-680 nm bandpass range for emission collection, was acquired. Twenty-five frames were acquired in high-speed mode (768 × 768 pixels) from each mouse retina and automatically co-registered and averaged in real-time using the Heidelberg Eye Explorer (HEYEX 1) software automatic real-time (ART) processing feature. Averaged images were auto-normalized for best contrast between hypo- and hyper-fluorescent/reflective features. Images were exported as TIFF files.

Following cSLO, SD-OCT images with a 45° FOV (∼1.4 mm) were collected with the optic nerve centrally positioned in the image FOV. Orthogonal B-scans (1400 A-scans/2 B-scans × 15 frames/B-scan) were collected at 0° and 90° to capture the horizontal and vertical meridians through the optic disk. Upon completion of the imaging, mice were administered 1.5 mg/kg of atipamezole HCL (Modern Veterinary Therapeutics, LLC., Miami, FL, USA) to induce recovery from general anesthesia. Eyes were covered with petrolatum-based eye ointment (Puralube Vet Ointment, Dechra Veterinary Products) to protect the cornea from desiccation during recovery.

Fifteen individual SD-OCT B-scan image frames from each orthogonal B-scan were co-registered and averaged using InVivoVue software v2.1 (Leica Microsystems). Mean outer nuclear layer (ONL) thickness was measured from each regional quadrant (superior, inferior, temporal and nasal) halfway from the optic disk to the edge of the image window using the straight-line measurement tool.

### Electroretinographic (ERG) Analysis

Mice were dark-adapted overnight for scotopic ERG testing and prepared as for *in vivo* retinal imaging. Following anesthesia, animals were placed on a stage maintained at 37°C. Custom-made clear plastic contact lenses with embedded platinum wires served as recording electrodes, and a platinum wire loop inserted into the animal’s mouth served as the reference electrode. ERGs were performed using an Espion E3 ColorDome ganzfeld apparatus (Diagnosys LLC., Lowell, MA, USA). For scotopic a- and b-wave responses, seven steps of increasing flash illuminance [-3.66 to −0.19 log candela (cd) s/m^2^] were presented in order of increasing flash strength. The number of successive trials averaged together decreased from 10 for the four lower-level flashes to 5 for the three highest intensity flashes. b-wave amplitude was measured from the a-wave minima to the b-wave maxima at ∼5 & ∼40 ms following flash stimulus onset, respectively. Upon completion of dark-adapted testing, a steady 30 cd/m^2^ adapting field was presented in the ganzfeld bowl. Following 7 minutes of light adaptation, cone ERGs were recorded using strobe-flash intensities of 0, 0.3 and 0.6 log cd·s/m^2^ on a steady 30 cd/m^2^ background. The number of successive trials averaged together decreased from 20 for the first two flashes to 15 for the highest flash. Cone b-wave amplitude was measured from the pre-stimulus baseline to the positive peak of the waveform following the flash stimulus (22).

### Retinoid Analysis

For quantitation of ocular retinoids, mouse eyes (1 eye per sample) were homogenized in PBS containing 100 mM 0-ethylhydroxylamine HCL and neutralized with 4N NaOH to pH 6.5 essentially as described (39, 40). Subsequently, 1 ml methanol was added and all-trans-retinol acetate was added as an internal standard. After solubilization with hexane and centrifugation the sample was dried under argon and dissolved in acetonitrile. The sample was then analyzed using a reverse phase column (CSJ C18 column, Waters) and a Waters Acquity UPLC system with monitoring at 320 and 360 nm and the use of gradients of water (A) and acetonitrile (B) containing 0.1% of formic acid as follows: 0-5 min, 60% B; 5 to 60 min, 60-70% B; 60 to 70 min, 70 to 100% B; and 70 to 90 min, 100% B min (flow rate of 0.3 ml/min). Absorbance peaks were identified by comparison to external standards.

### Plasma Lipid Analysis

Blood was collected directly from the heart using a heparin coated syringe and plasma was separated by centrifugation. Total plasma cholesterol (23-666-202) and triglyceride (23-666-412) concentrations were enzymatically measured using kits (Thermo Fisher Scientific) as described (25, 28).

### Statistical Analyses

All graphs and statistical analyses were performed using GraphPad Prism version 9.5.1. (Graph Pad, San Diego, CA). *t*-tests were performed between samples to determine p-values with an alpha-value of 0.05.Two-way ANOVA was performed when comparing more than 2 different variables. All data were analyzed and graphed as Mean ± SEM. A value of p ≤ 0.05 was considered significant and represented as * *p* < 0.05, ** *p* < 0.01, *** *p* < 0.001, *no asterisk: p* > 0.05.

## Results

### Absence of MTP expression in the RPE leads to decreased local Apo-B containing lipoprotein (Blp)

Blps are a major component of retinal degeneration-associated deposits (21). To test the hypothesis that the local assembly of lipids into Blps supports RPE and PR health and maintains visual function, we generated transgenic mice with an RPE-specific knockout of *Mttp*. To generate *RPEΔMttp* mice, *Mttp^flox/flox^* mice (with loxP sites flanking exon 5 and 6 of *Mttp*; <u>NM 008642.3</u>) were crossed with BEST1-Cre transgenic mice (24, 29). To confirm the excision of the *Mttp* gene in *RPEΔMttp* mice, we examined the expression of *Mttp* in the RPE by *in situ* hybridization. In control mice, *Mttp* BaseScope signal was detected in the RPE, while the RPE of *RPEΔMttp* mice produced little detectable signal (Fig. 1A). As expected, MTP protein (97kDa) was barely detectable in the RPE of *RPEΔMttp* mice compared to RPE of control mice (immunoblots show ∼80% decrease) (Fig. 1B). Consistent with studies in other cell types showing that MTP is required for the assembly of Blps, (25–28, 41), loss of RPE *Mttp* expression was associated with a decrease in Blp levels, as measured by APOB ELISA, in the RPE/Ch (Fig. 1C). APOB levels in the NR were not significantly different between *RPEΔMttp* mice and controls (Fig. 1C). In flat mounts immunostained for APOB, the RPE of *RPEΔMttp* mice showed ∼60% fewer APOB positive puncta compared to control mice (Fig. 1D). Of note, Blps can be delivered to the RPE via the choroidal circulation, which could partially explain decreased rather than complete loss of Blp associated with the RPE/Ch fraction. Collectively, our results confirm the ablation of Mttp expression and consequential decrease in Blps in the RPE of *RPEΔMttp* mice.

**Figure 1.**
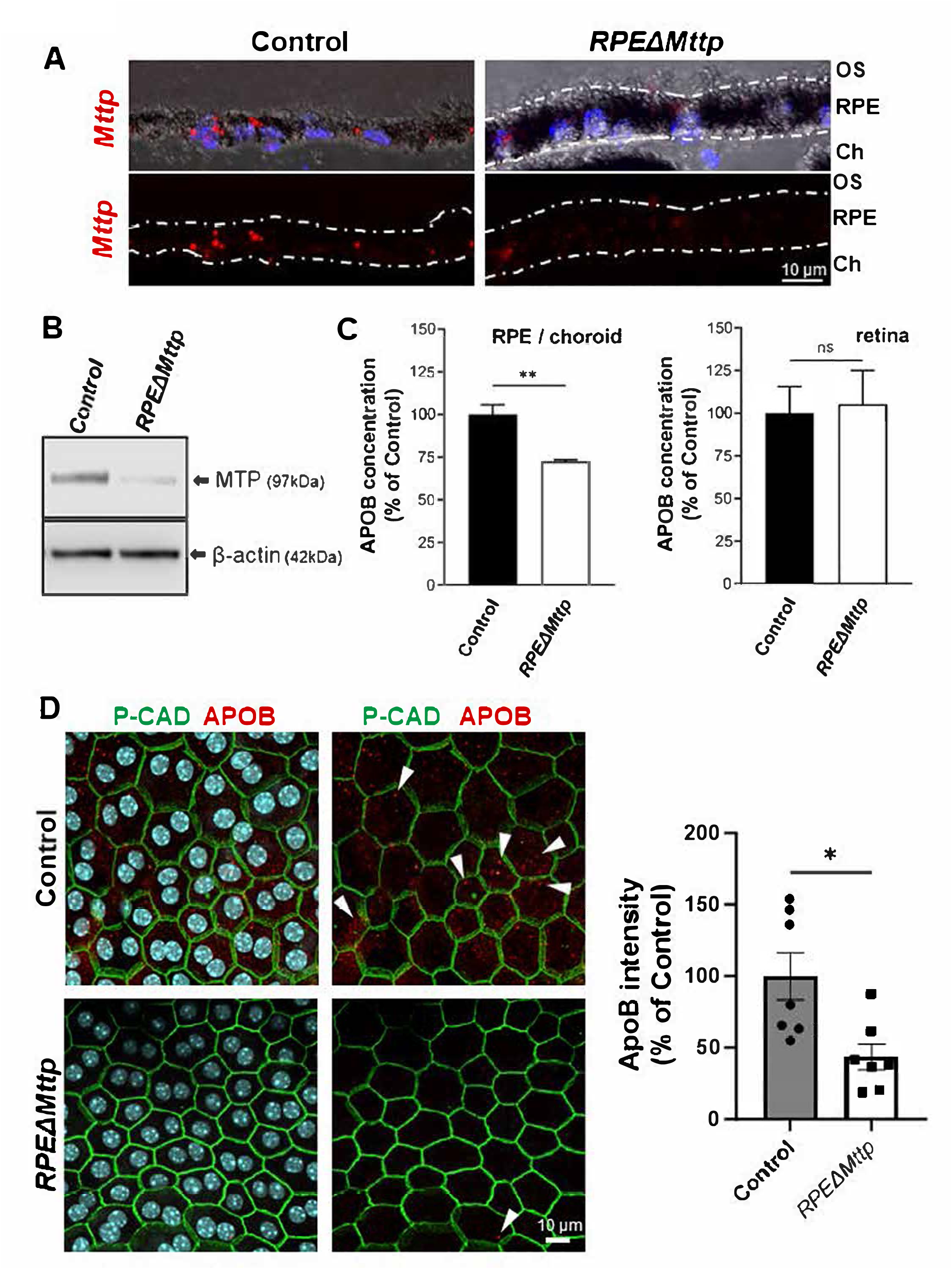
Ablation of *Mttp* in the RPE leads to decreased β-lipoproteins in the RPE. **A.** RNA *in situ* hybridization to *Mttp* expression in cryosections from 12-month-old *RPEΔMttp* mice and controls. Signal from the *Mttp* transcript (*red*) was detected in the RPE of control mice but diminished in the RPE of *RPEΔMttp* mice; Hoechst nuclear stain (*blue*), transmitted light (*upper images*); white dashed lines indicate apical and basal membranes of RPE layer. **B.** Immunoblot analysis of protein lysates from the RPE/choroid of *RPEΔMttp* mice and controls probed for MTP and normalized to β-actin. **C**. APOB decreases in *RPEΔMttp* RPE but not in *RPEΔMttp* retina. APOB concentrations in the RPE/choroid and NR of *RPEΔMttp* mice and controls were measured by ELISA in 9 month-old mice. Values shown are mean APOB concentration (± SEM) as a percentage of control in the RPE/Ch (*left*, t-test, *p* < 0.01) and neural retina (*right*, t-test, *p* > 0.05) normalized to total protein (*n* = 3). **D.** Representative immunofluorescence images showing RPE flat mounts from 4 month-old *RPEΔMttp* mice and controls stained for p-cadherin (CDH3, *green*) and APOB (*red*) with Hoechst nuclear stain (*cyan*); white arrows indicate APOB-stained particles. Mean APOB intensity per ROI (± SEM) was quantified (t-test, *p* < 0.05).

### RPE-specific MTP depletion has no effect on systemic lipids or most ocular retinoids

To determine if ablation of *Mttp* expression in the RPE altered systemic lipid concentrations, which may then affect RPE and PR health in *RPEΔMttp* mice, we measured cholesterol and TG levels in the blood plasma of *RPEΔMttp* mice and age-matched controls. Mean systemic cholesterol and TG concentrations were not significantly different between *RPEΔMttp* mice and controls (Fig. 2A). Quantification of ocular retinoids, showed no significant difference in total retinoid, all-trans-retinal (atRAL), all-trans-retinol (atROL), and 11-cis-retinal (11cisRAL),concentrations between *RPEΔMttp* and control eyes (Fig. 2B). In contrast, all-*trans*-retinyl ester (atRE), the storage form of retinoid was decreased by ∼30% in the *RPEΔMttp* eyes compared to control (Fig. 2C). atRE is associated with lipid droplet-like structures (42). These results suggest that ablation does not affect systemic lipid levels and the observed changes in the retina and RPE of *RPEΔMttp* mice are most likely due to the RPE-specific loss of MTP, rather than alterations to systemic lipid levels or vitamin A deficiency.

**Figure 2.**
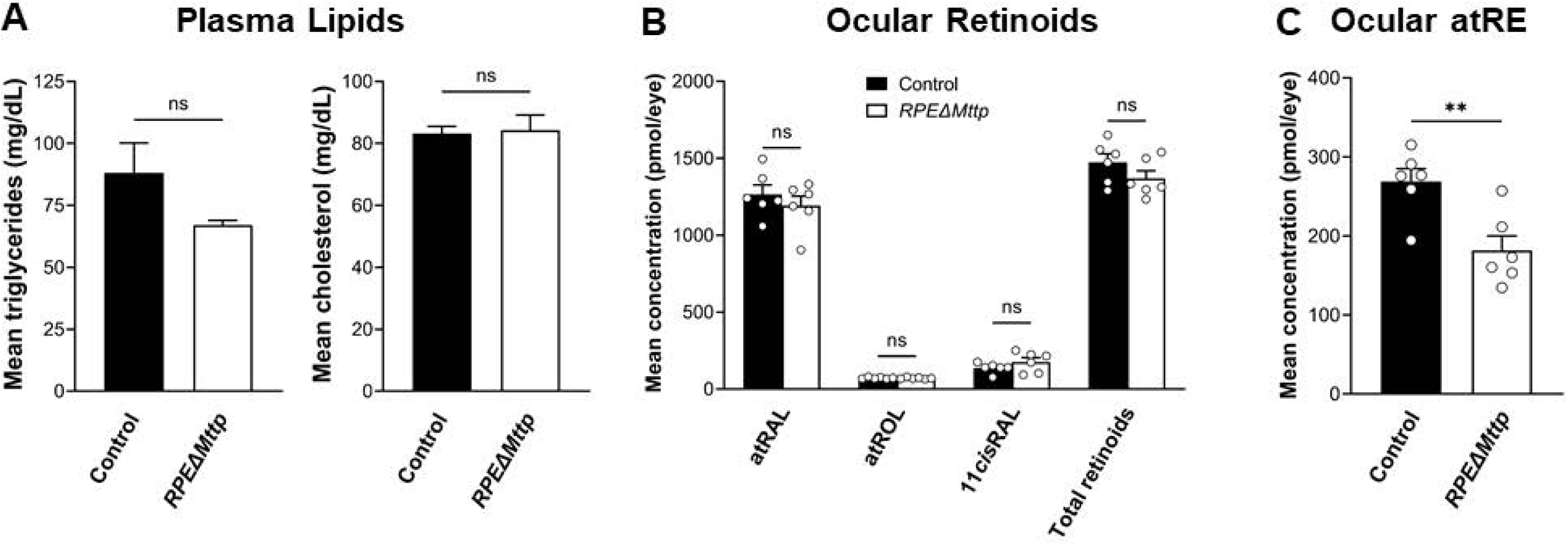
RPE-specific knockout of *Mttp* does not modulate plasma lipid but reduces ocular retinoid storage. **A.** Concentrations (± SD) of cholesterol (*left*, t-test, *p* > 0.05, ns) and triglycerides (*right*, t-test, *p* > 0.05) in blood plasma of 3-4-month-old were not significantly different between *RPEΔMttp* mice and controls (*n* = 4). Values are mean (± SEM). **B.** Mean (± SEM) all-*trans*-retinal (atRAL), all-*trans*-retinol (atROL), 11-*cis*-retinal (11*cis*RAL), and total retinoids concentrations in whole eyes from 3 - 4-month-old mice were not significant different between *RPEΔMttp* mice and controls (*n* = 3, two-way ANOVA, *p* > 0.05, ns). **C.** All-*trans*-retinyl ester (atRE) concentrations were decreased in *RPEΔMttp* mice. Mean atRE concentration in whole eyes from 3 - 4-month-old *RPEΔMttp* mice and controls (*n* = 3, t-test, *p* < 0.01)

### Retinal degenerative changes in mice lacking RPE-specific MTP expression

We next examined the impact of RPE-specific MTP depletion on retinal integrity in the *RPEΔMttp* eye using *in vivo* imaging (22, 43). Both blue autofluorescence-(BAF-) and infrared dark field-(IRDF-) cSLO images indicated a change in autofluorescence and reflective heterogeneity in 5- and 7-month-old *RPEΔMttp* mice compared to age-matched controls (Fig. 3A). Auto-fluorescence observed in *RPEΔMttp* mice was more spatially heterogeneous than in controls (Fig. 3A, *green arrows*). Such profound changes often originate from cellular alterations at the RPE level, giving rise to a mottled spatial appearance (22). Using SD-OCT, we monitored changes in the retina and show representative B-scans from the horizontal meridian for 5- and 7-month-old *RPEΔMttp* eyes and age-matched controls (Fig. 3B). We highlighted a region of interest just to the nasal side of the optic nerve (Fig. 3B, *gold box*) that contains a hyper-reflective feature indicative of intra-retinal pathology (*gold arrow*) in the 7-month-old *RPEΔMttp* mouse. This lesion appears to incorporate and IS and OS bands of photoreceptor cell attributable OCT lamina processes observed in clinical studies (44). Similar changes were not detected in 3.5-month-old *RPEΔMttp* mice (S.Fig.1). To determine if outer retinal abnormalities correlated with photoreceptor degeneration, ONL thickness was measured from the same SD-OCT B-scans. At 5-months of age, the ONL thickness of *RPEΔMttp* mice was reduced by over 1.5 µm relative to controls, with this difference doubling by 7 months of age (Fig. 3C).

**Figure 3.**
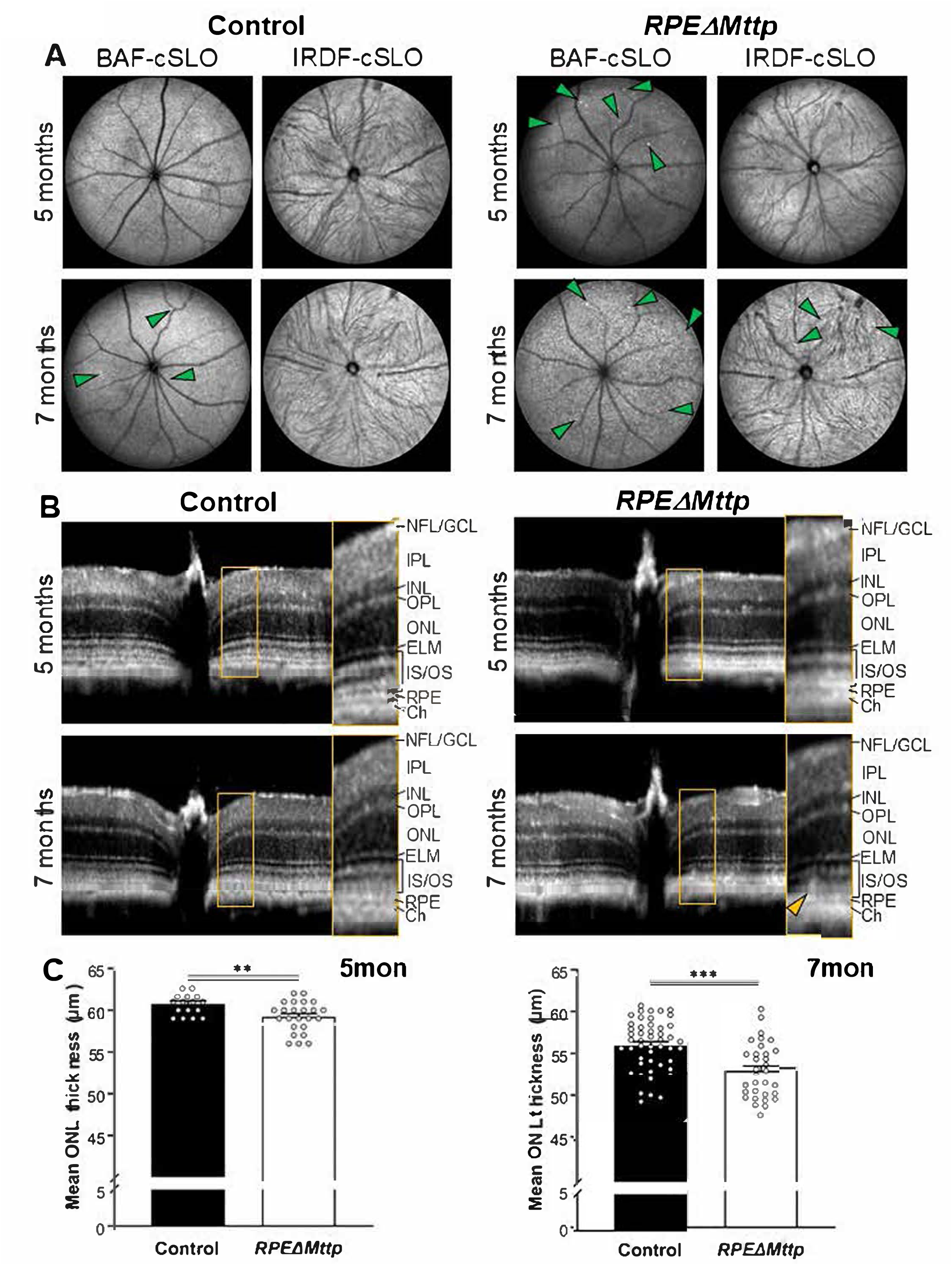
Photoreceptors degenerate in mice lacking RPE *Mttp* expression. A.–B. *In vivo* ocular imaging of 5 month-old and 7 month-old *RPEΔMttp* mice and controls. **A.** Representative blue autofluorescence-(BAF-) and infrared dark field-(IRDF-) cSLO images with hyper-autofluorescent or hyperreflective foci (*green arrows*), respectively **B.** Representative OCT images with a hyperreflective lesion visible in outer retina of the 7 month-old *RPEΔMttp* mouse (*gold arrow*). NFL/GCL: nerve fiber layer/ganglion cell layer, IPL: inner plexiform layer, INL: inner nuclear layer, OPL: outer plexiform layer, ELM: external limiting membrane, IS/OS: photoreceptor inner/outer segments **C.** ONL thickness decreased in *RPEΔMttp* mice. Mean ONL thickness (± SEM) measured from OCT scans of *RPEΔMttp* mice and controls at 5 months (t-test, p < 0.01) and 7 months (t-test, p < 0.001).

### Loss of MTP in the RPE leads to RPE lipid imbalance and dysregulation of retinal cholesterol

Collectively, the SLO and SDOCT imaging studies suggested alterations in RPE that may be indicative of lipid accumulation. Staining RPE flat mounts for neutral lipids with LipidTOX revealed strong lipid accumulation in RPE cells of *RPEΔMttp* mice (Fig. 4). The mean area of LipidTOX-positive structures (lipid aggregates) in the *RPEΔMttp* mice was 1.7 times that observed in age-matched controls. In control RPE, neutral lipid structures with an average area of 0.92 µm^2^ ±0.083 µm^2^, were observed, In comparison, in the *RPEΔMttp* mice, the average area of neutral lipid structures was1.587 µm^2^ ±0.182; a ∼ 45% increase in size.

**Figure 4.**
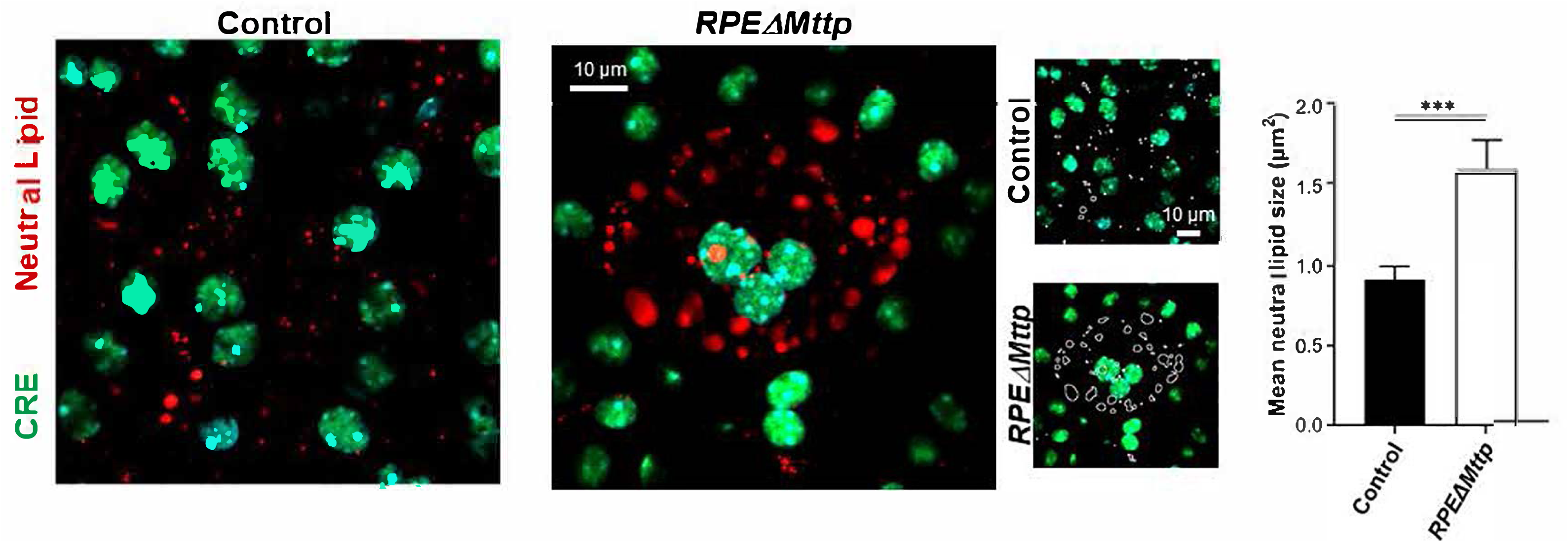
Lipid accumulates in the RPE and neural retina of *RPEΔMttp* mice. **A.** Neutral lipids accumulated in the RPE cells of *RPEΔMttp* mice. RPE flat mounts from 8-month-old *RPEΔMttp* mice and controls stained for Cre recombinase (*green*) and neutral lipids (LipidTOX, *red*). Mean area (± SEM, *p* < 0.001) of LipidTOX-positive aggregates (*outlines shown*) were quantified using FIJI.

MTP transfers cholesterol, triglycerides and phospholipids in the assembly of Blps, therefore we evaluated the effects of RPE-specific MTP depletion on cholesterol homeostasis in the retina by staining with filipin which binds to the 3’-OH group of unesterified cholesterol (UC). While filipin positive puncta were observed in the basal region of the RPE in controls, similar puncta were barely detected in the *RPEΔMttp* mice (Fig. 5A left panel, white arrows). Interestingly, intensely stained cholesterol puncta were detected within the outer segment and inner segment regions of *RPEΔMttp* mice compared with controls (Fig. 5B, right panel, yellow arrows). Numerous bright filipin foci within the ONL were also observed, similar foci were less abundant in controls (Fig. 5B). Filipin staining was ∼3-fold higher in the OS layer compared with control. Biochemical assessment of cholesterol levels in the NR of the *RPEΔMttp* mice, showed ∼ 35% increase in cholesterol relative to control (Fig. 5C). The increase in NR-associated cholesterol correlated with increased levels of RPE cholesterol transporter, ABCA1 (Fig. 5D); there was on average a 165 ± 18% increase in ABCA1 in *RPEΔMttp* compared to age matched controls. No change was detected in, the ATP-dependent cholesterol transporter, ABCG1 (102±0.76%) between *RPEΔMttp* and controls. These data suggest that in the absence of MTP activity, photoreceptor cells retain or accumulate more cholesterol in OS and photoreceptor cell-associated laminae. The change in ABCA1 implicates modulation of cholesterol efflux.

**Figure 5.**
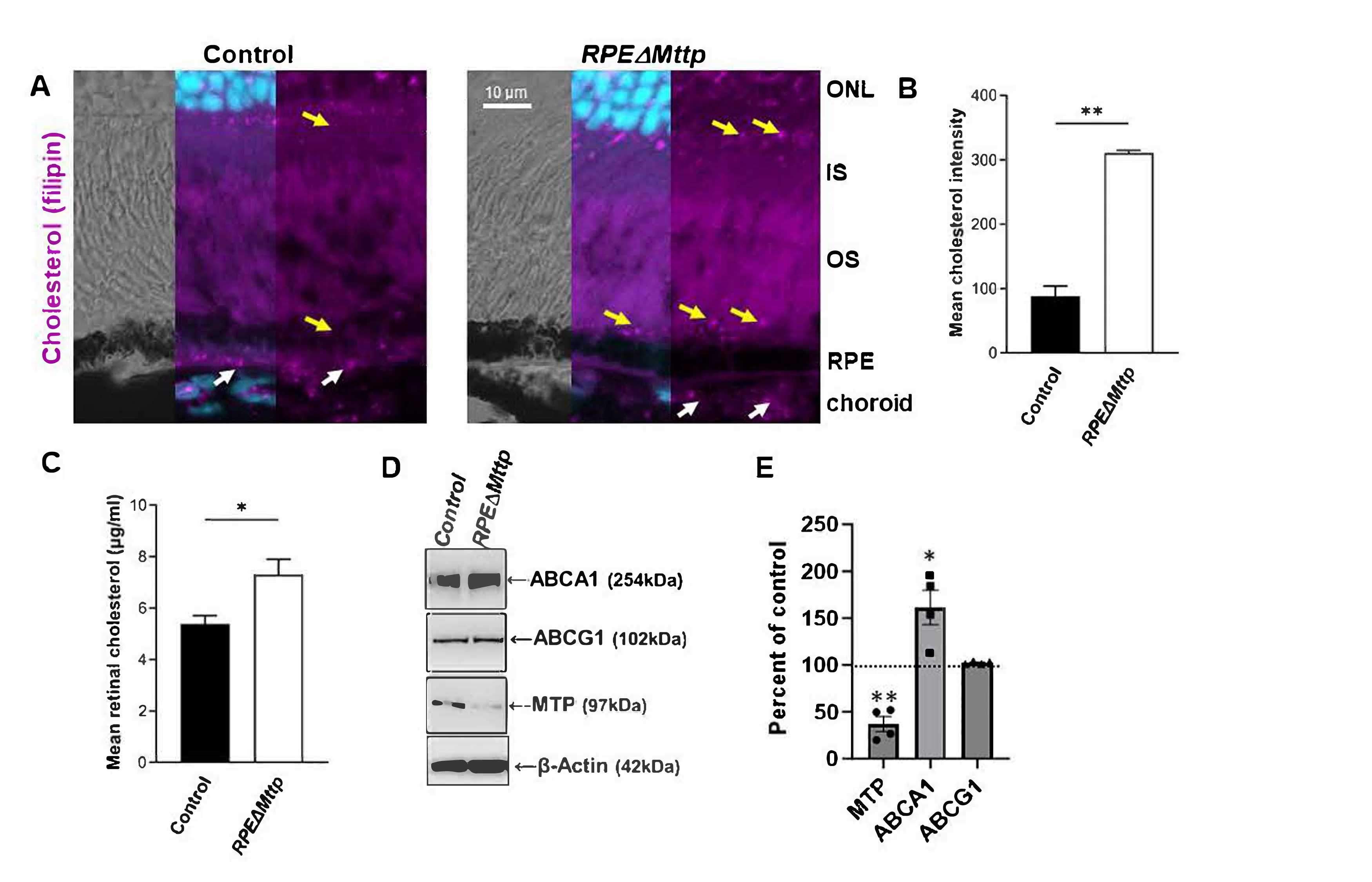
Unesterified cholesterol accumulates in the NR of *RPEΔMttp* mice. **A** Retinal cryosections from *RPEΔMttp* mice and controls probed for cholesterol (Filipin, *purple*). Mean UC (± SEM) intensity was quantified in the OS region (t-test, *p* < 0.01). **C.** Total cholesterol increased in the NR of *RPEΔMttp* mice. Mean retinal cholesterol concentration (± SEM) in the NR of *RPEΔMttp* mice and age-matched controls (t-test, *p* < 0.05).

### Structural and functional consequences of disrupted lipid homeostasis in the RPEΔMttp mice

*RPEΔMttp* mice exhibited retinal degenerative changes, as well as RPE and retinal lipid imbalance, thus we tested if the loss of RPE-MTP depletion affects retinal structure. Retinal cryosections from *RPEΔMttp* and control mice were stained for cone matrix sheaths with peanut agglutinin lectin and for rod photoreceptors with anti-opsin. The outer retina of *RPEΔMttp* mice routinely (all *RPEΔMttp* mice studied) exhibited sporadic structural alterations resembling tubulations (Fig. 6A and B), similar structures were not observed in any of the Ctrl mice studied. The regions around the *RPEΔMttp* dysplasia appeared to have shorter photoreceptor IS/OS thickness with relatively normal cone and rod density (Fig. 6C-6F). Photoreceptor degeneration is often accompanied by immune cell infiltration and microglial action in the subretinal space (45, 46). The BAF scans (Fig. 3A) reveled hyper-fluorescent foci in 5-month-old mice. To verify that the hyper fluorescent foci correspond to microglia, retina cyryosections were labeled with an antibody to Iba1. *RPEΔMttp* tubulations were often associated with Iba 1 positive cells in the sub-retinal space suggestive of inflammatory microenvironment in the affected areas (SFig.2). As previously described, autofluorescence associated with the microglia and detectable at 488 nm excitation likely derives from phagocytosed bisretinoid fluorophores that form in the photoreceptor outer segments populating the interiors of the tubulations (47).

**Figure 6.**
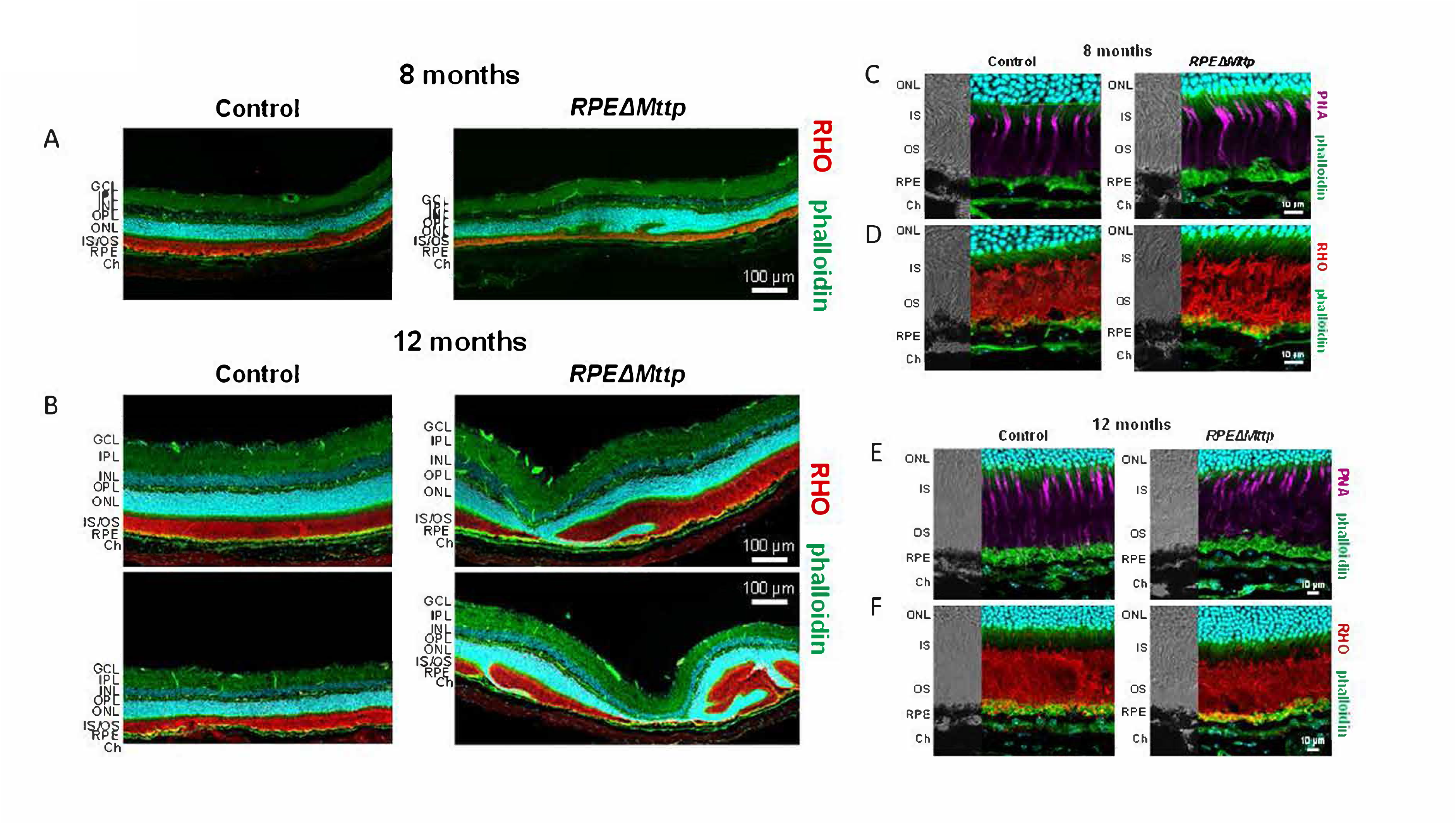
Structural anomalies in *RPEΔMttp* mice. Retinal cryosections from A. 8-month-old and B. 12-month-old *RPEΔMttp* and age-matched control mice stained with phalloidin (F-actin, green), Hoechst nuclear stain (cyan) and probed for opsin (RHO) (red). Retinal dysplasia was evident in *RPEΔMttp* mice but not age-matched controls. Retinal cryosections C and D, 8-month-old and E and F, 12-month-old *RPEΔMttp* and age-matched control mice stained with phalloidin (F-actin, green) and Hoechst nuclear stain (cyan) and either peanut agglutinin lectin (PNA, magenta) in Panels C and E or probed for opsin (RHO) (red), Panels D and F.

By 5 months of age, the retinas of *RPEΔMttp* mice showed morphological anomalies (Fig. 3 and Fig. 7). To characterize the nature of these anomalies, we examined retina ultrastructure using transmission electron microscopy. Electron microscopy showed an irregular RPE border and swollen RPE cells with disorganized basal infoldings (SFig. 3). Toluidine blue stained semi-thin sections the *RPEΔMttp* revealed hypertrophic and hyperpigmented RPE (Fig. 7A). This cell appeared to be full of pale spherical inclusions that are relatively large, homogeneous and electron lucent (Fig. 7A, Inset). As shown in Fig. 7B, we also observed large deposits located in the sub-retinal region consisting of various size inclusions mixed with disordered outer segment fragments (Fig. 7B Inset, yellow arrows). In the following panels (Fig. 7C and D), we describe a series of morphological anomalies seen within numerous *RPEΔMttp* mice analyzed. Degenerating photoreceptor debris is observed in Fig. 7C, with additional areas within the RPE containing vacuolar structures (blue arrows). In Fig. 7D, we show a clear displacement of RPE apical microvilli abutting what appears to be a deposit consisting of granular debris (Fig. 7D, Inset, red arrows). Using an OTAP post-fixation method, we detected lipid-droplets adjacent to what appears to be electron dense, possibly melanolipofuscin granules in the *RPEΔMttp* (Fig. 7E, Inset, white arrow).

**Figure 7.**
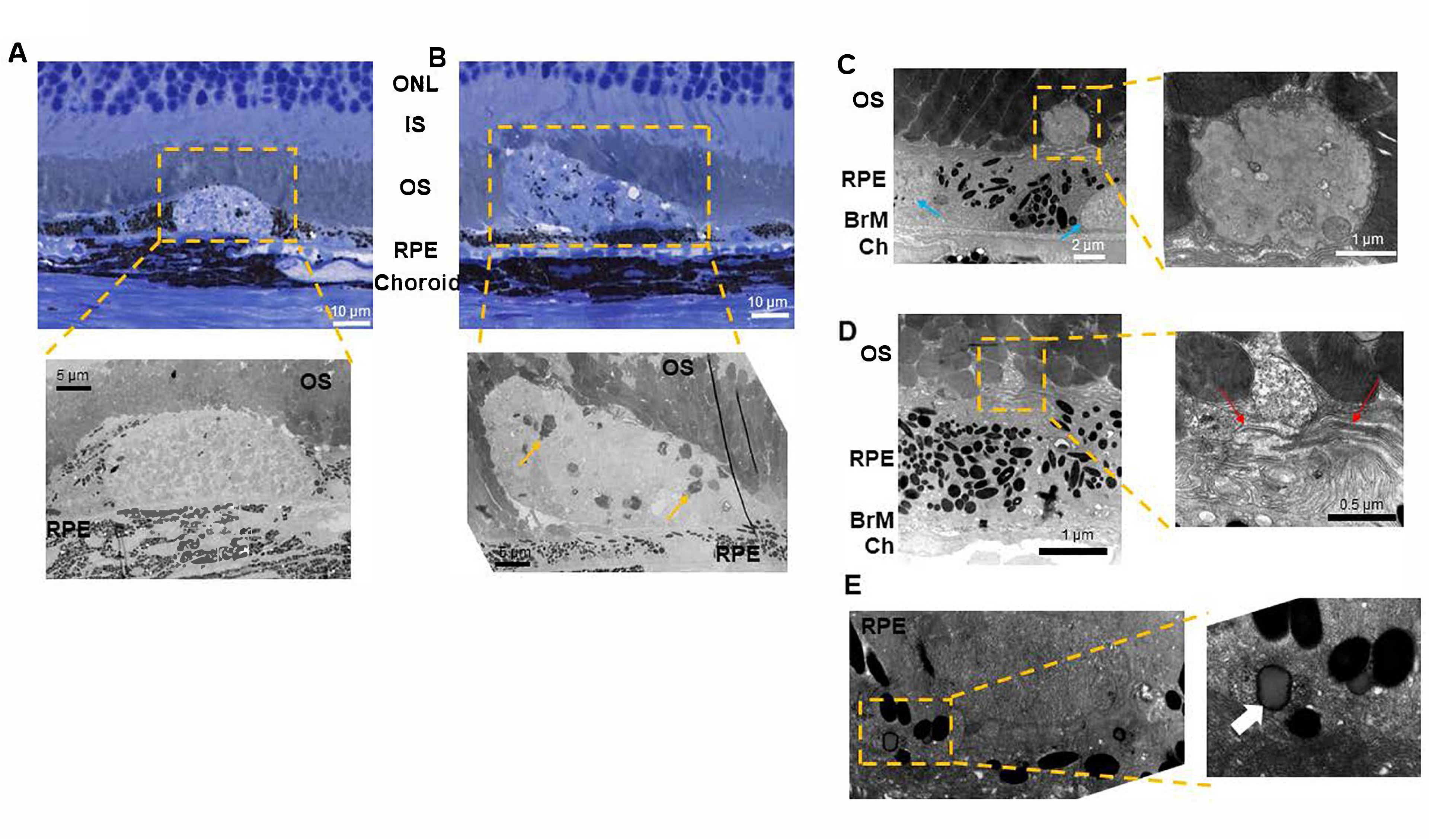
Morphological alterations in *RPEΔMttp* mice. TEM images of 8-9.5month-old *RPEΔMttp* mouse retinas. These images are representative of the types of anomalies we have observed in at least three *RPEΔMttp* mice. Orange boxes indicate areas enlarged to view detail. **A.** Toluidine blue stained semithin plastic section from 8-month-old, *RPEΔMttp* mouse eye centered on a swollen RPE cell. Inset, TEM, standard osmium fixation, shows this cell apparently full of spherical inclusions of varying sizes and electron density. **B.** Toluidine blue stained semithin plastic section from 8-month-old, *RPEΔMttp* mouse eye depicting subretinal deposit. Inset, TEM: standard osmium fixation showing enlarged debris, OS fragments are shown by the yellow arrows **C.** TEM using standard fixation of 9.5-month-old, *RPEΔMttp* mouse eye depicting retinal deposits, blue arrows indicate additional debris not shown in the enlargement. **D.** TEM using standard fixation of 8-month-old, *RPEΔMttp* mouse eye depicting deposit between RPE and PR cell layer. Red arrows indicate areas of apical RPE microvilli displacement, **E.** TEM using OTAP fixation of 9.5-month-old, *RPEΔMttp* mouse eye depicting lipid pool in RPE cell, indicated by white arrow.

### Visual Function

To gain insight into whether retinal lipid dysregulation impacts vision, luminance-response functions of the scotopic and photopic electroretinography (ERG) were compared between 5-month old *RPEΔMttp* mice and controls. Dark-adapted (rod photoreceptor driven) a-wave amplitudes were reduced by ∼50% of control in *RPEΔMttp* mice (Fig. 8A). Dark-adapted b-waves were reduced in a similar manner, likely due to diminished photoreceptor activation of bipolar neurons (Fig. 8B). No significant change in photopic b-wave was detected by 5 months (Fig. 8C). Taken together these results point to a functional decline associated with the rod photoreceptor pathway in the *RPEΔMttp* retina.

**Figure 8.**
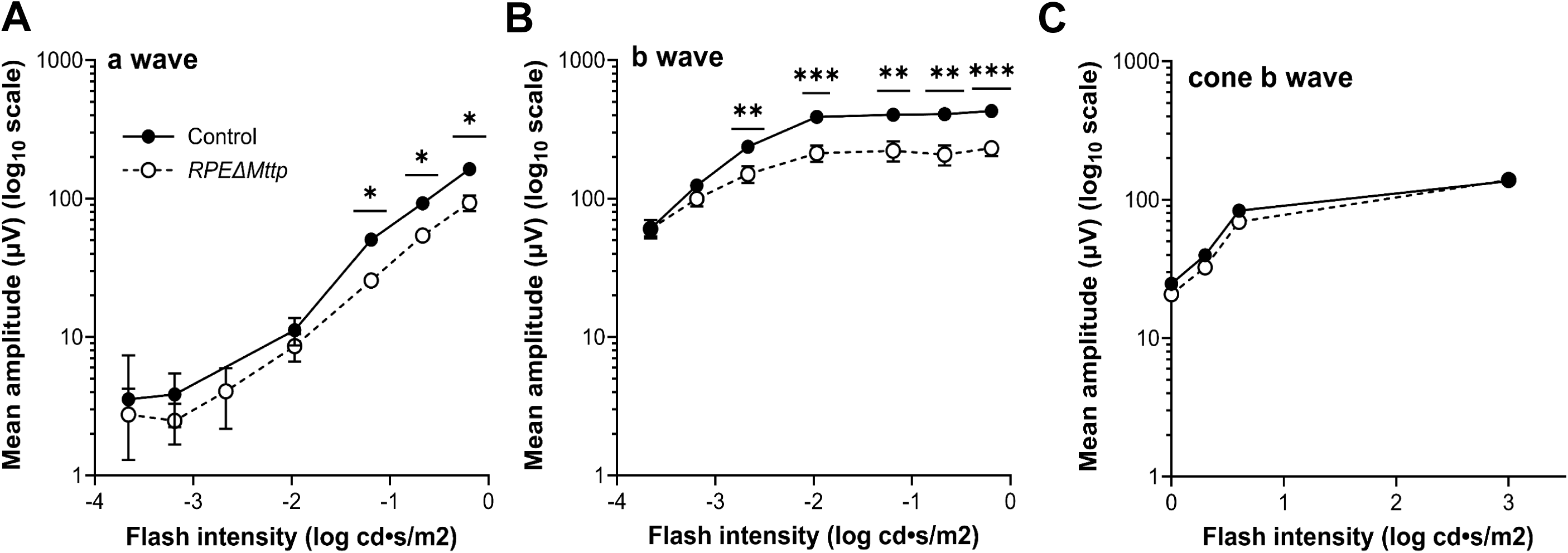
Loss of RPE-specific MTP decreases rod ERG response but does not affect cone function. Scotopic ERG a-wave (**A**) and b-wave (**B**) amplitudes and photopic ERG b-wave (cone b-wave) (**C**), plotted against flash luminance from 5-month-old dark adapted *RPEΔMttp* mice and controls. Data points indicate means (± SEM) of 3 *RPEΔMttp* mice and 3 controls, which were compared with t-tests (* *p* < 0.05, ** *p* < 0.01, *** *p* < 0.001, *no asterisk: p* > 0.05).

## Discussion

The ability of the RPE to maintain lipid balance and regulate the fate of its intracellular lipid pool is central not only to RPE but also photoreceptor function. The RPE intracellular lipid pool is fed by OS derived lipids through the daily ingestion of OS fragments by the RPE, in process historically referred to as phagocytosis but recently called trogocytosis (8, 48). We have retained the term phagocytosis herein. The daily phagocytosis and processing of polyunsaturated lipid rich structures is augmented with the *de novo* synthesis of lipids as well as the uptake of Blps from the systemic circulation as illustrated in Fig. 9. The studies herein point to RPE-MTP as an essential pathway in maintaining a healthy interplay between the RPE and photoreceptor lipid balance and hence cell function (for review (49))

**Figure 9.**
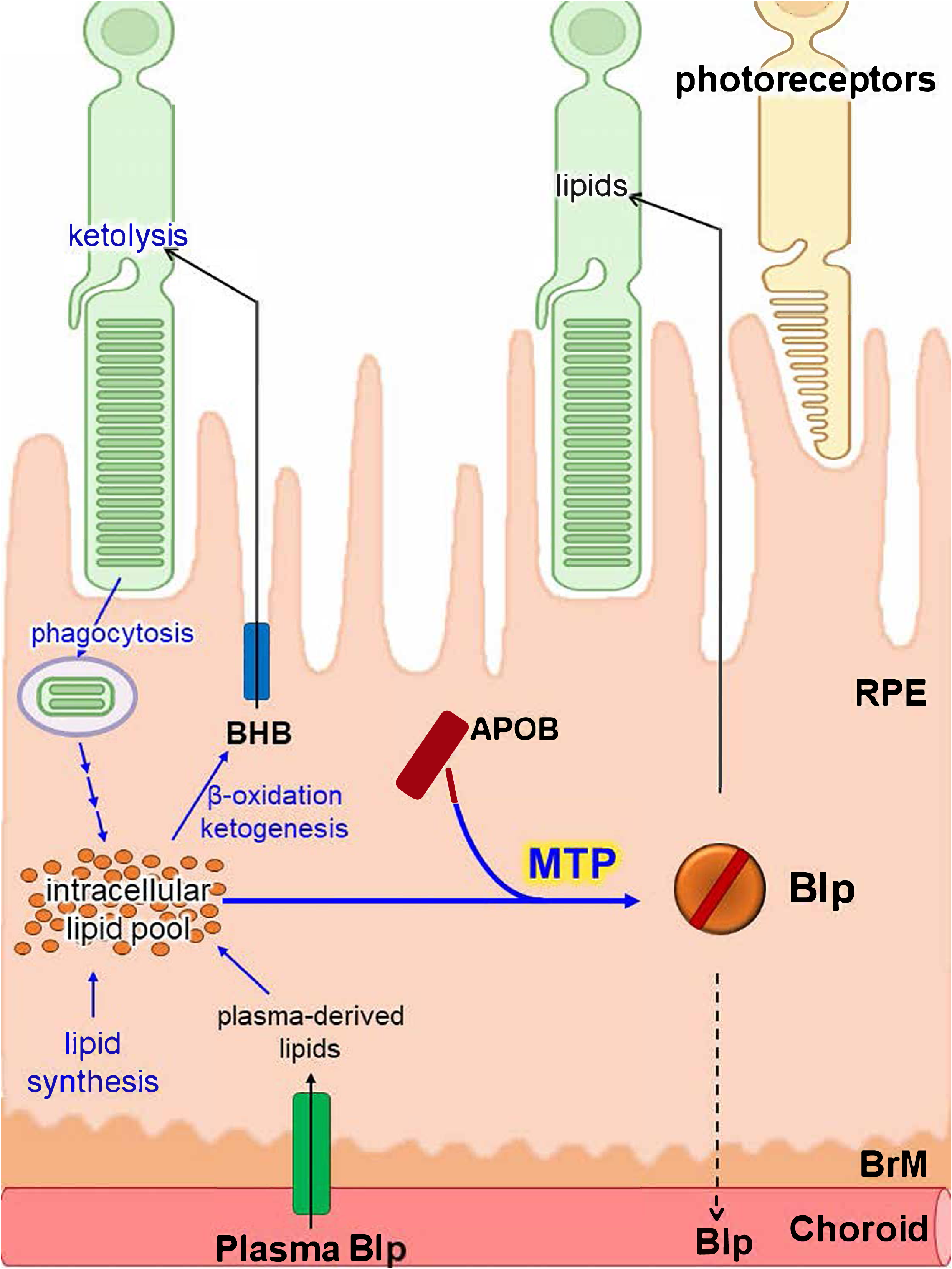
MTP activity is necessary to maintain RPE lipid homeostasis. Schematic representation of MTP-mediated Blp assembly in maintaining lipid balance in healthy RPE.

We generated and characterized the *RPEΔMttp* mouse model, in which the *Mttp* gene is deleted from the RPE to test the hypothesis that local Blps (i.e., those assembled in the RPE) are necessary to maintain retinal health and function. These studies fill a particularly understudied niche in retinal lipid metabolism. It was particularly important to develop a mouse model that could distinguish the role of local, RPE-derived Blps from that of systemic, plasma-derived Blps in retinal lipid homeostasis because intra-ocular Blps are distinct from those found in plasma. Compared to those found in plasma, intra-ocular Blps contain elevated levels of esterified cholesterol (EC; up 16- to 40-fold higher), carry lower levels of phosphatidylcholine, and display a different morphology (intra-ocular Blps are larger) (15, 21). In the *RPEΔMttp* mice, local Blp assembly was reduced while systemic lipids were unaffected. Consistent with MTP’s role in co-translational regulation of APOB expression, the RPE of *RPEΔMttp* mice contained less APOB than age-matched controls. As expected, RPE-specific knockout of the *Mttp* gene did not affect triglyceride or cholesterol concentrations in the blood plasma of *RPEΔMttp* mice, as these lipids are carried in systemic Blps that are mainly produced by the liver and intestine (17).

### RPE-MTP implications for systemic Blps and ABL

MTP deficiency in abetalipoproteinemia leads to progressive loss of vision. This is generally attributed to failure in the absorption of dietary fat-soluble vitamins. Besides the intestine, MTP expression has been shown in RPE. Here, for the first time, we have created a mouse model where systemic lipid metabolism is normal but is deficient in RPE MTP. This mouse model has normal plasma lipids. Despite normal systemic lipid levels, *RPEΔMttp* mice show significant AMD-like ocular abnormalities, such as retinal degeneration, hyper reflective foci, increased lipid deposition and reduced visual function as determined by ERG indicating that RPE MTP plays a crucial role retina lipid metabolism and some of the pathologies seen in abetalipoproteinemia patients might be related to RPE-specific MTP deficiency.

Abetalipoproteinemia (ABL) patients deficient in MTP activity have low plasma lipids and lipoproteins, as they are unable to synthesize chylomicrons and VLDL in the intestine and liver, respectively. ABL patients progressively lose eyesight with age (50, 51) with retinal dysfunction likely attributable to severe deficiency in the fat-soluble vitamins E and A. Vitamin A is necessary for phototransduction, while Vitamin E is neuroprotective with potent anti-oxidant properties, (6, 52). The highest levels of Vitamin E in the eye are found in RPE (52, 53); *in vitro*, RPE treated with Vitamin E have less complement activation (54). Major pathologic consequence of MTP deficiency include angoid streaks, retinitis pigmentosa and loss of vision with age (55–58), despite consumption of mega doses of fat-soluble vitamins (50, 51)

### RPE-MTP in retinal metabolic synergy

The retina relies on metabolic synergy for visual function; the RPE and retina have complementary metabolic roles such that they depend on each other for survival and individualized function. A critical aspect of this inter-dependence is the RPE’s metabolic flexibility. It relies on oxidative substrates; fatty acids, lactate and amino acids (proline) to provide TCA cycle intermediates thereby sparing glucose for the use by the neural retina (6, 59–62). Fatty acids are one of these flexible substrates, provided by the daily phagocytosis of lipid-rich OS, they generate fuel for RPE and NR function through fatty acid oxidation (FAO) (13). We would predict that RPE MTP-activity is necessary for oxidative metabolism to maintain RPE differentiation and function, which in turn regulates NR health.

### Depletion of RPE-MTP contributes to lipid accumulation in RPE and adjacent PR

The RPE shares metabolic similarity with cardiomyocytes; not only do both utilize fatty acids as an energy source (12, 63, 64), they both also secrete unique EC rich Blps (65–72). Several studies suggest that MTP-mediated secretion of Blps protects cardiomyocytes from lipid overload (73–76). iPSC derived cardiomyocytes from ABL patients fail to secrete apoB, accumulate intracellular lipids, and respond poorly to stress (77). Deficiency in RPE-MTP resulted in a similar intracellular accumulation of lipids. Given their particular lipid composition and role in intra-ocular lipid trafficking, RPE-derived Blps are proposed to play a role in the formation of drusen and other RPE-associated lipid accumulations (78). Drusen contain high concentrations of esterified cholesterol, unesterified cholesterol (UC), and TG but relatively little docosahexaenoic acid, a fatty acid abundant in PR, and also contain APOB, suggesting that drusen likely consist of RPE-processed material, rather than plasma-derived Blps or unprocessed phagocytosed OS (5, 20, 66, 71, 78). Subretinal drusenoid deposits (SDDs) that accumulate on the apical face of the RPE contain UC and the degraded remnants of shed OS (79, 80) but, unlike drusen, very little EC (80). While the origin of SDDs is not completely clear, dysregulation of lipid transport between the RPE and PR is implicated in their biogenesis (80).

Loss of RPE-specific MTP expression led to pathology in *RPEΔMttp* mice. *In vivo* imaging of 5-month-old *RPEΔMttp* mice revealed signs of retinal degeneration, such as hyper-reflective foci suggestive of lesions in the RPE, adjacent NR and ONL thinning, both of which were exacerbated with age. Lipids accumulated in the eyes of *RPEΔMttp* mice as neutral lipid aggregates detected in the RPE and cholesterol aggregates detected in the retina. Numerous models of dysfunctional lipid processing within the NR/RPE/choroid complex showed similar pathology to that observed here in the *RPEΔMttp*. Transgenic mice with ocular expression of a pathological mutant allele of human *ELOVL4*, an enzyme involved in the synthesis of very long-chain fatty acids, exhibited deterioration of structure and function in PR, steatosis and reduced OS processing, and eventual PR death (23). In the *Map1lc3b^-/-^* mice, encoding for LC3B, an autophagy-associated protein essential for efficient processing of phagocytosed OS, is deleted, leading to structural abnormalities in the NR and RPE, drusen-like basal lipid deposits, neutral lipid accumulation within the RPE, delayed processing of phagocytosed OS, and recruitment of immune cells (22). More specific to Blp dysfunction, two mouse models with the ablation of lipoprotein receptors that facilitate the internalization of Blp display pathology similar to that observed in the photoreceptors and BrM of AMD patients (81, 82). *Vldlr^−/−^* mice, which lack the receptor for very low-density lipoproteins (VLDL), display altered PR metabolism and PR secretion of the angiogenic protein VEGFA, leading to neovascularization and recruitment of Iba1^+^ cells (81). Lipids accumulate in the BrM of mice lacking the low-density lipoprotein (LDL) receptor (82).A recent bioinformatic analysis on dry AMD vs wet AMD identified *Mttp* as one of the top 41 most significant downregulated differentially expressed genes in dry AMD (83).

### Ocular retinoids and local (RPE-mediated) Blp assembly and secretion

The eyes of the *RPEΔMttp* mice in our study did not have lower total retinoid concentrations compared to age-matched controls. In contrast, loss of function mutations in *MTTP* as in abetalipoproteinemia (ABL) are associated with an absence of plasma Blps, leading to a decrease in the transport and availability of vitamin A and vitamin E to peripheral tissues, including the eye (5, 68, 71, 84). ABL patients show significant ophthalmologic complications (55–58) that are managed with vitamin A and vitamin E supplementation (85–87). Abetalipoproteinemia (ABL) patients deficient in MTP activity have low plasma lipids and lipoproteins and progressive vision loss with age (50, 51). Dietary, fat-soluble vitamin supplementation only delays the onset of vision loss, implicating a role for intestine specific MTP-activity in vision, a hypothesis that can be tested using conditional intestine specific MTP-deficient mice, *ENTΔMttp^IND^.* Blps also serve as systemic carriers of dietary vitamins, with Vitamin E serving as a potent anti-oxidant. Plasma Blps are required to transport fat-soluble vitamins out of the intestine and to the liver for storage, vitamin A is transported from the liver to eye via retinoid binding protein 4 (RBP4) (39). Although in the absence of *Rbp4*, RPE find a way to acquire retinol (vitamin A) by non-Rbp4 mechanisms that could involve lipoprotein. (39).Our results support the Blp--independent transport of vitamin A within the eye and suggest that the phenotype we observed is not due to vitamin A deficiency.

In contrast to the other retinoids quantified in this study, atRE was significantly lower in *RPEΔMttp* mice than in controls. In the RPE, retinoids are stored as retinyl esters in retinosomes, particles similar to intracellular lipid droplets (42). Given that MTP also regulates lipid droplet formation (88 34) we are testing, the hypothesis the MTP activity is necessary for retinyl ester storage in the RPE.

## Conclusion

In this study, we demonstrate that RPE-specific MTP activity is critical to maintain intracellular RPE lipid pools, likely by serving as a conduit to prevent lipid steatosis. When we deleted *Mttp* from the RPE we also observed cholesterol accumulation in photoreceptors and decreased rod cell function. On a molecular level, the accumulation of outer segment cholesterol is predicted to slow the plasma membrane delimited enzyme interactions governing phototransduction (89, 90) as is a decrease in DHA (91), the latter of which we predict is depleted due to decreased apcial RPE-Blp secretion. Cone cell function appears to be less dependent on MTP-mediated lipid processing. These studies have far-reaching implications for understanding how MTP expression and activity is regulated (92)in disease processes as diverse as iron-accumulation induced lipotoxicity of lysosomes (93), to the dysregulation of aerobic glycolysis-lipid OX-PHOS metabolic synergy in retinitis pigmentosa (94, 95) as well as diseases of the aging retina (5, 60). Moreover, value of this model is that the regulation of the RPE-Blp pathway can now be studied *in vivo*. This feature is of particular importance, due to the unique features of ocular Blps relative to other better known Blps from liver, intestine, and heart, and the singular feature of lipid handling due to OS phagocytosis by the RPE. Collectively, these studies strongly suggest that lipoprotein assembly, a key pathway for AMD drusen, is essential for outer retinal health.

## Abbreviation

11cisRAL: 11-*cis*-retinal
ABL: abetalipoproteinemia
APOB: apolipoprotein B
atRAL: all-*trans*-retinal
atRE: all-*trans*-retinyl ester
atROL: all-*trans*-retinol
Blp: apolipoprotein B-containing lipoproteins
Ch: choroid
cSLO: confocal scanning laser ophthalmoscopy
EC: esterified cholesterol
ERG: electroretinogram
IHC: immunohistochemistry
IS: inner segment
NR: neural retina
ONL: outer nuclear layer
OS: outer segment
PR: photoreceptor cell
RPE: retinal pigment epithelial cell
SD-OCT: spectral domain optical coherence tomography
TG: triglycerides
UC: unesterified cholesterol
VLDL: very low-density lipoprotein

## Acknowledgements

Grant support NEI core grant P30:EY001583, NEI EY0323743(KBB and MMH), NIHLB HL160470 and HL166214 (MMH). The donors of Macular Degeneration Research, a program of the BrightFocus Foundation, in the form of a Postdoctoral Fellowship (M2023002F) to CRG

## Data Availability Statement

Included in article. The data that supports the findings of this study are available in the Methods and or supplementary material of this article.

## Conflict of interest statements

The authors have no conflicts of interest

## Author contributions

Catharina Grubaugh, Anuradha Dhingra, M. Mahmood Hussain and Kathleen Boesze-Battaglia all contributed to writing of this manuscript. All authors contributed to editing the manuscript. Kathleen Boesze-Battaglia and M. Mahmood Hussain conceived the project. Kathleen Boesze-Battaglia, Catharina Grubaugh, Anuradha Dhingra, Lauren L. Daniele, Janet R. Sparrow and Mahmood Hussain designed the experiments. Kathleen Boesze-Battaglia, Catharina Grubaugh, Lauren L. Daniele and Anuradha Dhingra generated the *RPEΔMttp* mice. Binu Prakash performed the plasma lipid analysis. Catharina Grubaugh, Anuradha Dhingra, Lauren L. Daniele performed and analyzed all confocal imaging, morphology, biochemical and molecular assays. Brent A. Bell acquired and analyzed ERG data and performed and acquired SD-OCT and cSLO data. Janet Sparrow and Diego Montenegro performed the ocular retinoid analysis and provided expertise in Vitamin A delivery and distribution in retina. Christine A Curcio provided expertise in the interpretation of retinal histology.

## Notes

### Competing Interest Statement

The authors have declared no competing interest.

