## Supplemental Figures for "Microsomal triglyceride transfer protein is necessary to maintain lipid homeostasis and retinal function"

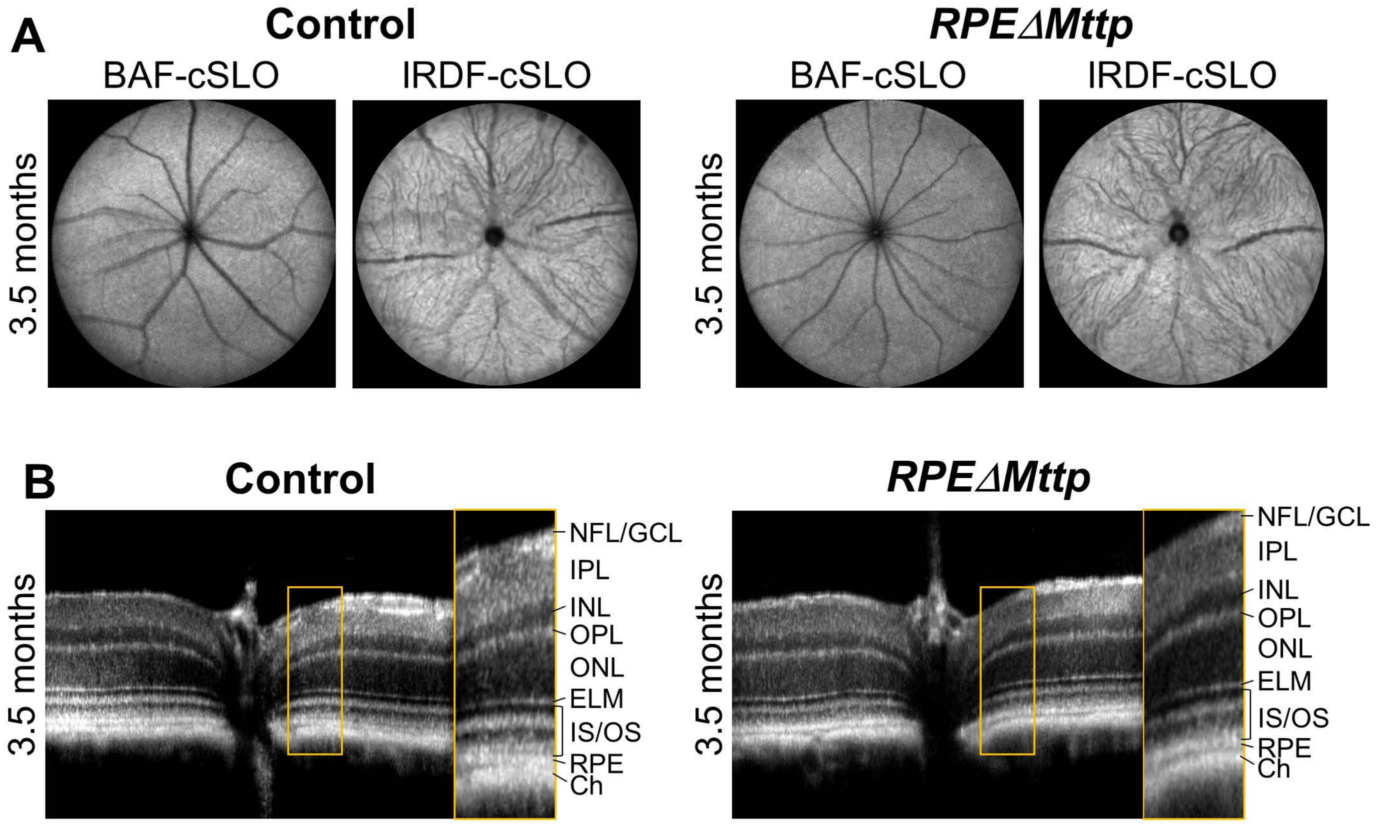


S Figure 1. In vivo ocular imaging suggest no change in 3.5month old RPEΔ*Mttp* mice. A.–B. In vivo ocular imaging of 3.5 month-old RPEΔ*Mttp* mice and controls. A. Representative blue autofluorescence- (BAF-) and infrared dark field- (IRDF-) cSLO images B. Representative OCT images of the 3.5 month-old RPEΔ*Mttp* and age-matched control mice. NFL/GCL: nerve fiber layer/ganglion cell layer, IPL: inner plexiform layer, INL: inner nuclear layer, OPL: outer plexiform layer, ELM: external limiting membrane, IS/OS: photoreceptor inner/outer segments

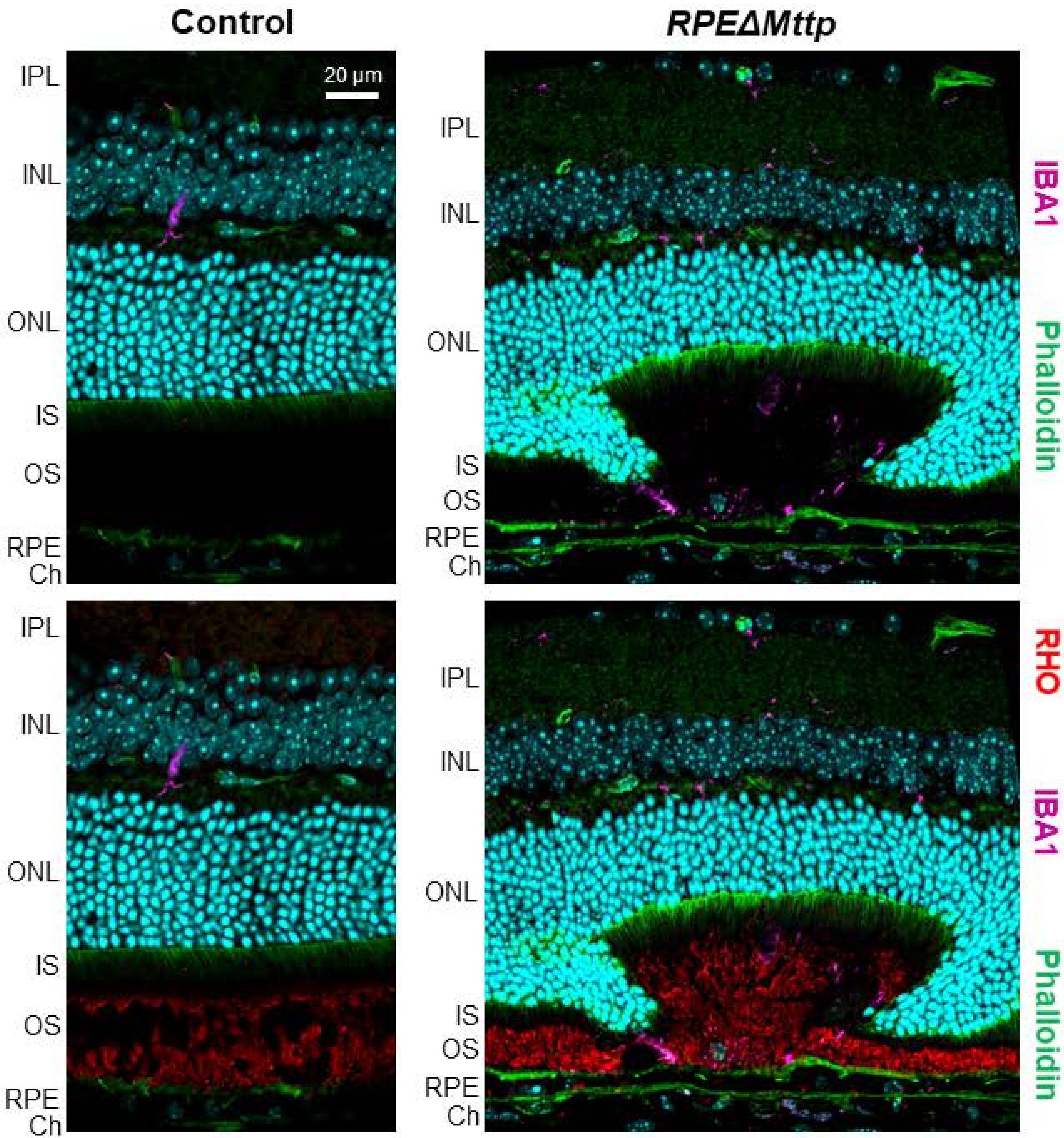


S Figure 2. Inflammation accompanies photoreceptor degeneration in RPEΔ*Mttp* mice. A representative rosette structure in the retina of the RPEΔ*Mttp* mouse is associated with Iba1 (magenta) staining; rhodopsin (red), phalloidin (green), Hoechst nuclear stain (cyan).

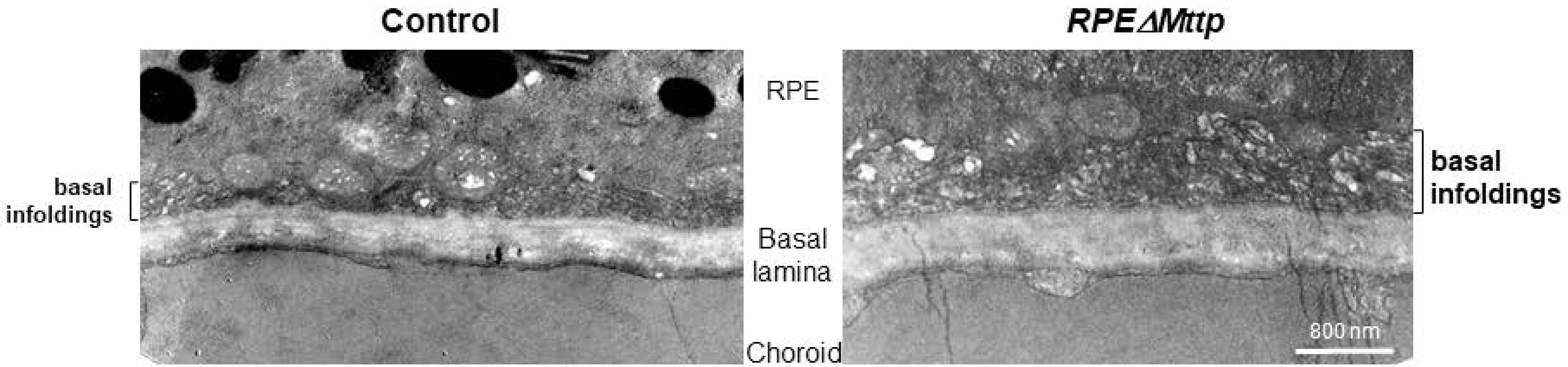


S Figure 3. RPEΔ*Mttp* mice exhibit disrupted basal infoldings. TEM using OTAP fixation of 9.5-month-old, RPE Δ Mttp and Ctrl mouse eye depicting basal infolding.
